# Harmonising distributed tree inventory datasets across India can fill critical gaps in tropical ecology

**DOI:** 10.1101/2024.05.18.594774

**Authors:** Krishna Anujan, Neha Mohanbabu, Abhishek Gopal, Akshay Surendra, Aparna Krishnan, Ankitha Jayanth, Tanaya Nair, Shasank Ongole, Mahesh Sankaran

## Abstract

1. Global analyses of tree diversity and function are strongly biased geographically, with poor representation from forests of the Indian subcontinent. Even though data from India - representing two-thirds of the subcontinent and spanning a wide range of tree-based biomes - exists, a barrier to syntheses is the absence of accessible and standardised data. Further, with increasing human footprint across ecosystems, data from Indian landscapes, with their long history of human-nature interactions is a key link to understand the future of tropical forested landscapes. Given the long history of human-nature interactions and high human footprint in the region, accessible and standardized data from the subcontinent can enable understanding the future of tropical landscapes under increasing human footprints globally.
2. Combining literature searches with manual data retrieval, we assembled the INdia Tree Inventory dataset, INvenTree. INvenTree is the largest meta-dataset of peer-reviewed publications (n = 465) from 1991-2023 on geolocated plot-based tree inventories of multispecies communities from Indian ecosystems, in aggregate covering 4653.64 ha and all of its woody biomes.
3. Using INvenTree, we show extensive sampling across tropical moist and dry forests, the dominant ecosystem types in the country. However, most studies have low sampling effort (median sampled area = 2 ha) and data across studies is not openly accessible (73.33 % of studies representing 83.43% of the sampled area), potentially hindering inclusion into regional or global syntheses.
4. Significantly, we show majority authorship from within the country; 82.8% of corresponding authors were from India and 73.33% of the studies had all authors affiliated with Indian institutions. We also identify ecological and conservation sampling priority regions based on forest cover and forest loss and set a blueprint for future sampling efforts in the country.
5. Based on extensive Indian scholarship in forest ecology showcased through the INvenTree dataset, we see opportunity for regional collaboration to create scientific inferences that are larger and scalable, while prioritising data and knowledge equity. Harmonising these existing datasets, and synthesising historic and grey literature, will contribute enormously to understanding the human dimension of tropical ecology as well as informing regional management and conservation.

## Introduction

Distributed tree inventories, typically collected using standardised methodologies, are powerful tools for addressing fundamental macro- and micro-ecological questions. Tree inventory information is a crucial baseline to understand ecological communities and their susceptibility and resilience in the face of global change including advancing our understanding of forest structure and dynamics, carbon stocks and fluxes, and improving predictions under climate change scenarios (Anderson-Teixeira et al., 2015, 2015, 2018; Cazzolla Gatti et al., 2022; ForestPlots.net et al., 2021, 2021; Jung et al., 2021; Malhi et al., 2002; Poulter et al., 2019). However, the global distribution of these tree inventory datasets is geographically biased, leading to highly variable and weakly constrained predictions from data-poor regions, many of which overlap deforestation and climate-change hotspots (Carvalho et al., 2023).

Continental biases in data pose a barrier to truly global ecology, both because of ecological and anthropic differences among the global tropics. Firstly, tropical biomes show continental scale differences in species richness, stem density and carbon stocks, because of differences in earth history and biogeography (ForestPlots.net et al., 2021; Hagen et al., 2021; Sullivan et al., 2017). Secondly, there is growing recognition of historical and ongoing human influence in shaping natural landscapes, influencing patterns, processes and functioning (Barlow et al., 2012; Fraser et al., 2024). This human influence on ecosystems has differences across continents and biomes, with some tropical biomes having only <1% under low human influence (Riggio et al., 2020; Venter et al., 2016). Yet, global analyses using tree inventories draw on sparse and biased data from the tropics, hindering truly “global” ecological knowledge (Martin et al., 2012; Stocks et al., 2008). Global syntheses use publicly available data (e.g. hosted at GBIF or Dryad) and aggregate national scale data (e.g. US Forest Inventory and Analysis), inadvertently reflecting and propagating the biases within these datasets. At the global scale, collection and compilation of biodiversity data is biased towards countries with high domestic product, high proportions of English speakers and high levels of security (Amano & Sutherland, 2013; Hughes et al., 2021). Temperate biomes are oversampled, while tropical biomes are understudied, and the Neotropics are studied more than the Paleotropics (Martin et al., 2012; Stocks et al., 2008). For a robust understanding of tropical ecology in the Anthropocene, ecological understanding must span variation on both ecological and human dimensions.

The Indian subcontinent is a pronounced gap in most global datasets and analyses, occluding a robust understanding of both ecological and human dimension of tropical ecology. The Indian subcontinent is one of the oldest and most biodiverse regions of tropical Asia, with a long human history and present-day footprint. Human settlement here has been recorded as early as 700,000 to 400,000 years before present (Briggs, 2003; Gadgil & Thapar, 1990; Mani, 1974). Among the global tropics, this region ranks highest in present-day human footprint on ecosystems, on par with highly built areas of the industrialised temperate regions (Venter et al., 2016). To this day, these ecosystems continue to support high human densities and human use, for timber, fuelwood, fodder and non-timber forest products (Karanth & DeFries, 2010; Shahabuddin & Prasad, 2004-04/2004-06). Despite this unique combination of bioclimatic, biogeographic and human forces in shaping present-day natural landscapes, the subcontinent continues to be excluded from macroecological syntheses, including estimates of tree density, forest age and species richness (e.g. Cazzolla Gatti et al., 2022; Crowther et al., 2015; Poulter et al., 2019). These geographic gaps could be due to a combination of factors including gaps in data collection within the subcontinent (e.g. across biomes or regions of interest) or gaps in data reusability (e.g. data stored in repositories are missing data or metadata components).

Within the Indian subcontinent, data availability and accessibility could be affected not only by sampling constraints, but by legacies of scientific inequalities, further hindering collaboration and synthesis. Stemming from colonial legacies, tropical ecology research and publishing has been has been dominated by researchers from high-income countries, leading to “parachute science” (Maas et al., 2021; Miller et al., 2023; Nuñez et al., 2021; Odeny & Bosurgi, 2022; Raja et al., 2021; Stocks et al., 2008). Moreover, despite ecological research increasingly moving to a big-data and collaborative discipline, its costs and benefits continue to be unevenly distributed (Farley et al., 2018; Maas et al., 2021). This is especially stark for synthesis and meta-analyses requiring quantitative skills and no new data collections (Mori et al., 2015; Stocks et al., 2008). As voiced by researchers in the Neotropics and in Africa, researchers based in the Global South experience multistep challenges in the scientific publishing pipeline - lack of funding, language barriers, inequalities at the peer-review stage and inability to pay high open access publishing costs - as well as transient and transactional partnerships with Global North researchers (Asase et al., 2022; Kwon, 2022; Mekonnen et al., 2021; Ramírez-Castañeda, 2020; Smith et al., 2023). The Indian subcontinent, with legacies of its recent colonial past reflected in society, including in ecology and forestry practices, is likely affected by these same issues. A large and diverse region such as the Indian subcontinent is prone to parachute science by Global North researchers into the region, as well as parachute science within the region: researchers with more resources may extract data and knowledge from distant, often resource-poor regions within the subcontinent. Such academic inequalities could disincentivise data sharing, especially in the absence of clear incentives (De Lima et al., 2022). Since barriers to data access have strong social dimensions, it is important to carefully examine the geographic distribution of the researchers, biomes and publication patterns, to understand the community of data owners and assess collaboration needs.

As the largest country in the Indian subcontinent, and representative of its diversity and human footprint, data collation from India can further ecological understanding for the region, beyond biodiversity assessment. India, currently the most populous country in the world, continues to have large forest cover and high species diversity under four biodiversity hotspots (Mittermeier et al., 2004; Mugal et al., 2023; Myers et al., 2000). Ecosystems in India cover 10 biomes as proposed by Dinerstein et al. (2017), covering all biomes of South Asia. Moreover, ecosystems in India are human-dominated; they continue to support high human densities along with high biodiversity to the extent that 85% of high priority conservation areas in India are outside of protected areas and embedded in wider human-nature landscapes (Karanth & DeFries, 2010; Srivathsa et al., 2023). India, therefore, provides a unique “model” to understand human-dominated tropical landscapes as the human footprint on natural landscapes across the world continues to grow (Karanth & DeFries, 2010; Roberts et al., 2021). Further, Indian scholarship in forest ecology is extensive, with a growing number of research programmes and investment, including over 250 long-term monitoring efforts (Tiruvaimozhi et al., 2024). Given its large human capital, collaborative frameworks from India can also set benchmarks for inclusive tropical ecology. Here, we assembled INvenTree, the largest meta-dataset of geolocated plot-based tree inventories from India and assessed (i) sampling patterns, (ii) geographic gaps in sampling and (ii) authorship patterns in tree community research in India to aid future syntheses.

## Materials and Methods

### Study area

The Indian subcontinent is one of the oldest regions in tropical Asia (Briggs, 2003; Mani, 1974). Its complex geo-climatic and biogeographic history has resulted in a rich variety of biomes with high diversity and endemism of flora and fauna (Fig 1). Within the subcontinent, India, 6° 440 and 35° 300 N latitude and 68° 70 and 97° 250 E longitude, comprises the largest landmass, with four global biodiversity hotspots occurring within mainland India (Mittermeier et al., 2004; Myers et al., 2000). It comprises ten biogeographic subdivisions based on phytogeography (Rodgers & Panwar, 1988), and five major broad forest types are commonly recognised: tropical moist forest, tropical dry forest, subtropical forest, temperate and sub-alpine forest (Champion & Seth, 1968; Reddy et al., 2015) (Fig S1). Among these, the tropical dry forest is the most dominant vegetation type in terms of land cover (Reddy et al., 2015). The current distribution of flora within the mainland is predominantly shaped by the interplay of past and current climatic events and biogeography (Mani, 1974). Seasonal rainfall driven by the Himalayan orogeny is one of the major drivers of current vegetation patterns with most of the subcontinent strongly governed by monsoonal rainfall. For ease of comparison with global studies we used the classification proposed by Dinerstein et al. (2017), which resulted in classification of the Indian mainland and the Andaman and Nicobar Islands into ten biomes with several ecoregions nested under these (Fig 1, Fig S2).

**Figure 1:**
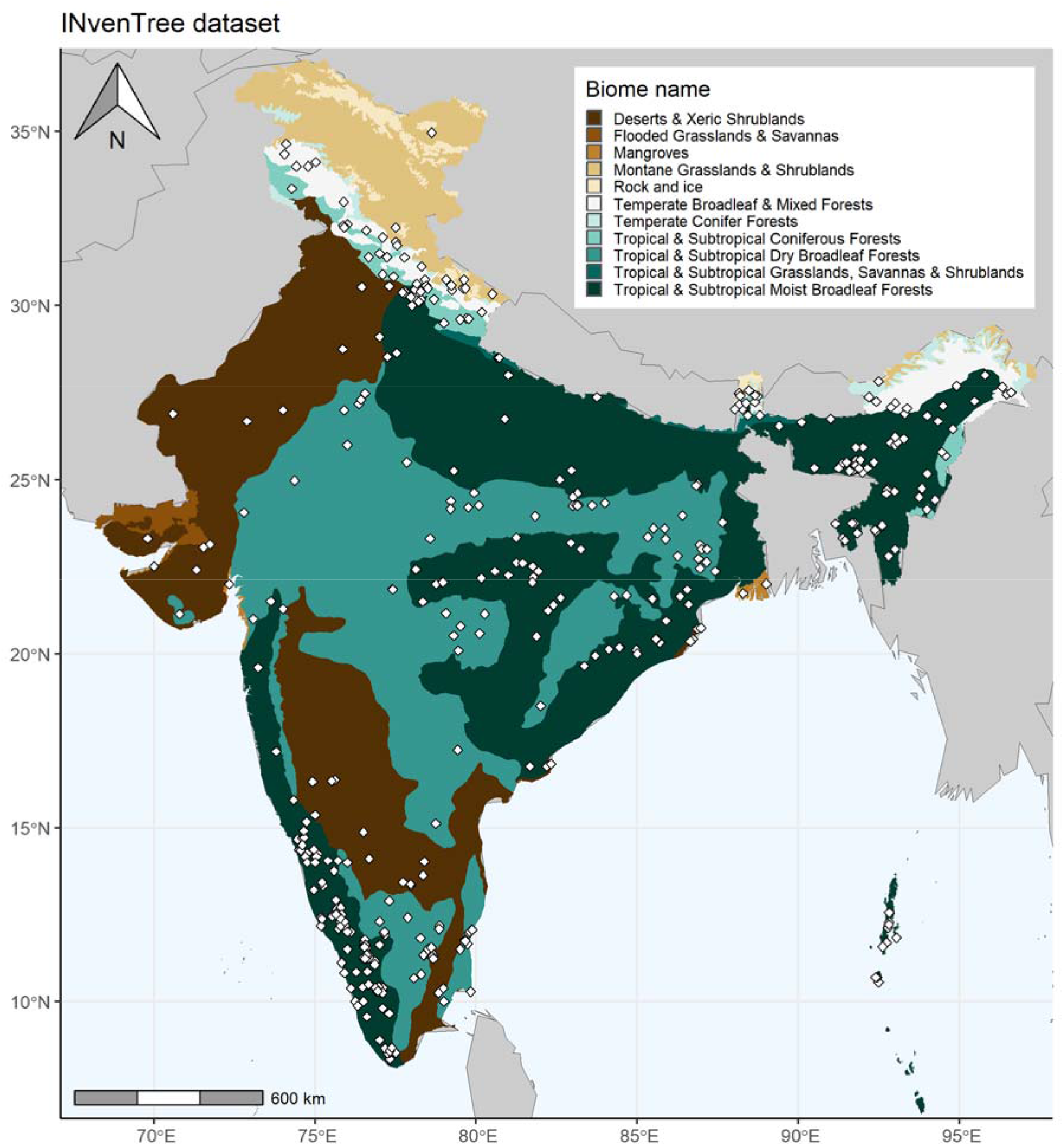
INvenTree dataset. The location of study sites in papers that form part of the INvenTree dataset in relation to the 10 vegetated biomes (and Rock and Ice) in India as per the Dinerstein et al. (2017) dataset. Each point corresponds to the centroid of the sampled location reported in each study.

The Indian subcontinent has a long history of human populations with the earliest human settlement recorded between 700,000 to 400,000 years before present (Gadgil & Thapar, 1990). Human population spread throughout the subcontinent at low densities in the terminal pleistocene (20000 - 10000 years BP) and by 4000 years BP, agricultural settlements had covered the entire subcontinent. India is currently the most populous country in the world, with a population of 1.4 billion which is projected to continue rising in the coming decades [World Population Prospects]. To this day, forested landscapes in India also house large densities of human populations with forest- or other natural-resource-dependent livelihoods.

### Assembling the INvenTree dataset

We created a meta-dataset of plot-based tree inventory studies on multispecies communities from Indian forested ecosystems from 1991 to 2023. We used both Google Scholar (last search in November 2021) and Web of Science (last search in February 2023) to include a wide range of journals as well as book chapters that publish forestry research from India. The Google Scholar search terms included (Tree diversity OR plot OR dbh) AND (India) and was later updated to exclude other countries where “India” or “Indians” may have been used in other contexts (e.g.,other countries in the Indian sub-continent such as Pakistan, Nepal, Myanmar etc., American Indian reservations). We used a broader list of search terms on Web of Science with the keywords AB=((tree diversity OR forest structure OR (tree* AND biomass) OR (forest* AND biomass) OR carbon stock OR vegetation survey OR vegetation sampling OR (tree* AND plot) OR (forest AND plot) OR (measur* AND tree) OR (checklist AND flora) OR (checklist AND tree) OR (checklist AND plant*)) AND India). We supplemented this list with an additional set of papers we were aware of but weren’t present in the Google Scholar or Web of Science results (e.g. global studies that report on datasets from India but are not captured by the keywords), and with papers that we found through snowball sampling the known papers’ references. We merged the lists from all three sources and removed duplicates, resulting in 3758 entries. We then manually sorted the list based on the title and the abstract to remove papers that were not related to forests in India, resulting in 657 potentially relevant papers. We accessed shortlisted papers and screened them based on the following criteria– (1) the study involved collection of plot level or checklist data of trees, (2) the study sites were natural multi-species stands of trees from inland woody ecosystems or coastal forests and mangroves. We excluded papers focussing on seedlings, single species studies and those based in managed or urban environments. Based on these criteria we identified **465 papers that formed the INvenTree dataset**.

From each paper in the INvenTree dataset, we extracted data on i) geographic location of study ii) area sampled iii) author affiliations and iv) data accessibility. i) We recorded the latitude and longitude of the study area reported either from the text or from a map provided. For studies with more than one plot, where separate locations were reported for each, we estimated the centroid of the study area. ii) To calculate the total area sampled in each study, we recorded plot shape, plot dimensions and number of plots, for all plot-based studies. Since we were focussed on exhaustive inventories, for studies that sampled subplots within a larger plot (e.g. five 20 m x 20 m plots sampled within a 1 ha area), we recorded the total area of the subplots which were exhaustively sampled (5x 400 m^2^) instead of the reported study area (1 ha). Further, for studies that inventoried plots across natural and managed communities or plantations, we only counted plots that were in natural multi-species communities. (iii) We verified (for Web of Science search) or manually added information on author affiliations and addresses. (iv) To assess the accessibility of the full paper and data, we recorded whether the papers were freely available (open access) on the publisher’s webpage and whether individual level or species level summaries of the tree data were freely accessible in the paper, in appendices, or in open access data sharing websites like FigShare, Dryad or Zenodo. Finally, we identified papers that used the same datasets - either large scale datasets or long-term datasets. We excluded large-scale datasets from analysis of biome-distributions as they covered several biomes; for long-term datasets we included only the first published paper. For analyses of authorship, we used all published papers including large-scale and repeat datasets.

### Analysing data accessibility and reusability

We first examined the number of studies, total area and percent area sampled within each biome. We also examined the open access status and the data access status of studies across biomes. Although some of our sites fell outside of biomes expected to have tree cover (e.g. in Montane grasslands and shrublands), we included them if they included tree inventories, as this information is likely to be more accurate than biome classifications. To further understand the practical constraints on the reuse of these data, we assessed each dataset on two axes: the data resolution and reusability. We considered these axes independently because for full reusability for synthesis, data should be available at the finest resolution, stem level data, but some synthesis might be possible with aggregate data as well. For each study that had reported data beyond broad summaries, we accessed the data from external repositories or from the paper.

We used the FAIR framework for scientific data management (Wilkinson et al., 2016) to assess the reusability status of dataset in INvenTree, by categorising datasets in each study based on whether they are Findable, Accessible, Interoperable and Reusable. We categorised these datasets on the FAIR-ness axis, for three different data resolutions - plot-level, species-level and stem-level data. We defined plot-level data as data aggregated at the plot level, where plot is the smallest spatial unit of sampling, species-level data as data aggregated at the species level, whether within or across plots, and stem-level data as data reported at the level of data collection - individual stems. For detailed methods, refer to the supplementary material.

### Analysing geographic patterns in sampling

We assessed sampling gaps in India in two different ways; we compared the spread of our data in relation to a) tree cover (as a proxy of ecological sampling priority) and b) its loss in India (conservation sampling priority). Indian forested ecosystems have a large range of natural tree cover, spanning closed canopy forests to open natural ecosystems. For robust ecological understanding, sampling should span this range of variation in tree cover. We defined ecological sampling priority as undersampled areas across the tree cover gradient. We also defined conservation priority sampling areas as areas of high tree cover loss but low sampling effort. We used the Hansen et al. (2013) dataset of tree cover in the year 2000 (within the period of interest) and tree cover loss from 2000-2021 cropped to the limits of India using Google Earth Engine (code here - https://code.earthengine.google.com/1f887c3af348300241dfc60e3da21cff). For these 30 m x 30 m pixels, we followed Friedl & Sulla-Menashe (2022)’s definition of forested pixels as >30% forest cover. This would include woody savannas (30-60% cover) but exclude more open and grassy savannas (10-30%).

India has 594 districts as of the 2011 census with a median area of 4077.48 km^2^ ranging from 15.7 km^2^ to 47858 km^2^ area. We chose district as our sampling unit as the sampling scale of most of the studies in the INvenTree dataset was contained within a district. Moreover, ecological heterogeneity is often small enough within a district and summaries at this scale are meaningful for further use by multiple stakeholders (Srivathsa et al., 2023). We used an administrative map of districts from [GADM] and extracted the tree cover (number of forested pixels) and cover loss (number of pixels experiencing loss) for each district using the *extract* function from the *terra* package (Hijmans, 2022b). For each district, we then calculated our variables of interest - % tree cover (percentage of district area with tree cover) and % forest loss (number of pixels that experienced forest loss divided by number of forested pixels). We chose to analyse % forest loss instead of absolute forest loss since there is a large distribution of forest cover across districts and absolute loss would be somewhat correlated to forest cover (for analysis of absolute forest loss, refer to the supplementary appendix). We then summarised the proportion of area sampled in each district as the sum of the total area sampled in each study that fell within its boundaries divided by the district area. We limited the analysis to districts that have non-zero forest cover (n=341) and excluded all points that fell outside these even though they may have reported on tree-based biomes, because of high uncertainties.

To analyse the spatial distribution of data, we assigned each district into one of three categories (low, medium, high) along each variable assessed - % tree cover, % forest loss and sampling effort. For %tree cover, we split districts as three quantiles and scored them as 1-3, i.e. districts that fell in the lowest 1/3rd of % tree cover values was scored as 1. Many districts had zero tree cover loss (n=115 out of 341) or zero sampling effort (n=205) and so, it was not possible to split the districts into three evenly sized groups based on the values of these variables. Moreover, zero sampling and zero loss are qualitatively different from low sampling and need to be highlighted. We, therefore, assigned all districts with zero tree cover loss as category 1, and split the remaining districts evenly between categories 2 and 3. We used a similar logic for sampling effort, but since our aim was to highlight research gaps, for sampled effort, we assigned a reverse scale. All districts with no sampling were categorised as the highest “sampling gap” and assigned category 3. The remaining districts were split evenly between categories 1 and 2. In our scale, a district falling in the highest quantile of tree cover and no sampling effort would be scored (3,3) to highlight high priority ecological sampling. Similarly, a district with high loss and no sampling effort would be scored (3,3) to highlight high priority conservation sampling. We visualised spatial overlaps in tree cover or tree cover loss and sampling gaps using bivariate chloropleth maps. We repeated these steps for sampling gaps across ecoregions. For detailed methods and data distributions, refer to the supplementary material.

### Analysing author affiliations and parachute science

We considered the full INvenTree dataset (n = 465) for the bibliometric analysis. For each unique study, we extracted the addresses of all authors. For corresponding authors, we used the address provided through Web of Science or extracted from the paper manually. Some publications in our dataset did not have an identifiable corresponding author, in which case it was left empty. In cases where the corresponding author had multiple affiliations, we considered only the first. For each study, we identified the unique affiliation addresses and the number of authors affiliated to each address. We ran all unique addresses from papers through a Google Maps-based geocoding software on Google Sheets, Awesome Table Geocode, that returns latitude and longitude for each address (https://awesome-table.com).

To analyse collaboration potential within the region for future syntheses, we compared the geographic positions of the authors and the papers in different ways. For these analyses, we limited the dataset to only those studies where we had information on the location of the authors as well as the site (n=456). First, we examined the proportion of authors in each study that were affiliated with institutions in India and its relation to the area sampled. Second, we examined the distance of the study site from the corresponding author institution for each study, by calculating the Haversine distance (the shortest distance between the two points assuming a spherical earth) between the two pairs of coordinates using the *geosphere* package (Hijmans, 2022a). We then looked at how this distance has changed over the years.

All analyses were performed in R version 4.1.3 (R Core Team, 2022) using packages *dplyr* and *tidyverse* (Wickham et al., 2019, 2022).

## Results

### Data availability and accessibility

The 465 studies reported in the journal articles in the INvenTree dataset span 10 biomes and cover 31 of the 36 states and union territories of India (Fig 1). We identified 436 studies of this dataset that report on area-based inventories, e.g. transects or plot-based measures, sampling a total of 4653.64 ha (Fig 1). The remaining were checklists, duplicates or did not report key metrics to estimate sampling effort. The most commonly studied biome was Tropical & Subtropical Moist Broadleaf Forests (236 out of 465, 50.75%), followed by Tropical & Subtropical Dry Broadleaf Forests (86 out of 465, 18.49%) (Fig 2a). In terms of area, the most extensively sampled biome was Tropical & Subtropical Moist Broadleaf Forests with a total area of 2169.3 ha sampled across 221 studies (Fig 2b).

**Figure 2:**
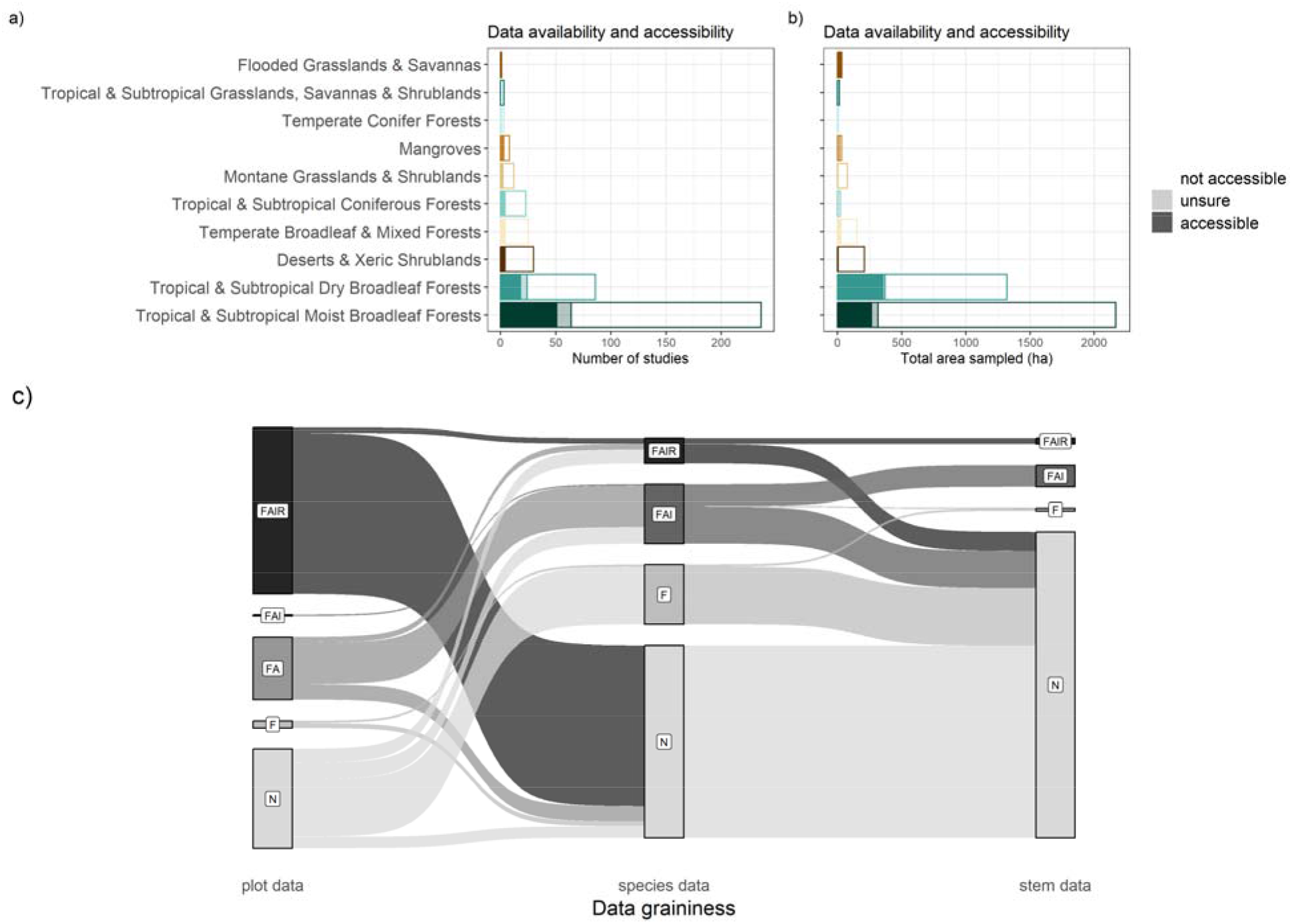
Data availability and accessibility across biomes. a) Number of studies and b)total area sampled across the 10 vegetated biomes of India among tree inventories studies in India. Solid bars represent data accessible in repositories or in the appendix while empty bars represent inaccessible data. Lightly coloured bars represent studies where the accessibility status was uncertain. Studies focussing on a single biome are only reported here. See supplementary material for details. c) Data distribution along data graininess and Findable, Accessible, Interoperable, Reusable (FAIR) principles of data reuse. Data reported only for studies where data was accessible for at least one of the categories of data graininess, the rest has been excluded from this figure. Legends F, FA, FAI, FAIR represent position along the reusability scale as defined by these principles. N represents data that were not findable. Size of the bars represent total sampled area in each of these categories. For detailed methods, refer to the supplementary material.

Most studies in the INvenTree dataset had low effort in terms of total area sampled; the median sampled area of the studies was 2 ha. Of these, 84.73% report on datasets that sample less than a total area of 100 ha, 72.47% less than 10 ha, 27.96% less than 1 ha. Across these studies, data accessibility was poor; only 101 studies, representing 21.72% of the studies and 14.8% of the area had data that was available either as a table, appendix, supplementary information or archived in a repository. Sampled area was not statistically significantly different between data-accessible and data-inaccessible studies (one-way ANOVA, df=1, SS=0.22, p=0.45). Data access was fairly evenly limited across the bioregions studied (-squared test statistic for number of studies = 40, p=0.1, for area = 80, p=0.2) (Fig 2b).

While examining the reusability of the datasets under the FAIR framework, we found considerable variation in the level of detail of findable and reusable data across studies that had “accessible” data (Fig 2c). Much of the datasets classified broadly as “accessible” (14.8% of the total sampled area) met reusability standards only for plot-level (12.81% of total sampled area across studies) or species-level data (1.95% of sampled area). Among the studies assessed, only 5 representing 0.47% of sampled area met reusability standards for stem level data, the finest grain of tree inventory information assessed. Despite limited data accessibility, the text of 56.77% of published articles were openly accessible (Fig S4). Studies were most commonly published in the journal Tropical Ecology (n=40) (Fig S4).

### Authorship patterns

Over 4 in 5 of the corresponding authors of papers in the INvenTree dataset were based in institutions in India (82.8%), with a small proportion of corresponding authors (14.19%) from other countries, primarily the United States (4.73%), France (2.37%) and China (1.72%) (Fig 3). Within India, affiliations were spread widely across the country, showing authorship and data ownership across institutions (Fig 3a). The number of publications have grown over the years and much of this increase is attributable to corresponding authors based in India (Fig S13). A clear majority, 1612 out of a total 1870 authors across articles in the meta-dataset were affiliated with Indian institutions and organisations (‘domestic’ in Fig 3), but 24.95% of papers also had authors affiliated with non-Indian institutions (‘foreign’ in Fig 3). When scored for the percentage of Indian-affiliated authors in a study, 90.11% of studies had more than 50% authors affiliated to India. 73.33% studies had all authors affiliated to Indian institutions. Although large-scale studies were few, they tended to have high or near-total in-country authorship (Fig 3). We observed an increasing trend of large-scale studies over the years (S13), but local and small-scale studies also continue to be published (Fig 3). Further, most studies in the INvenTree dataset were conducted locally, close to the institution of the corresponding author; the median distance between corresponding author affiliation and study site was 280.38 km (Fig 3).

**Figure 3:**
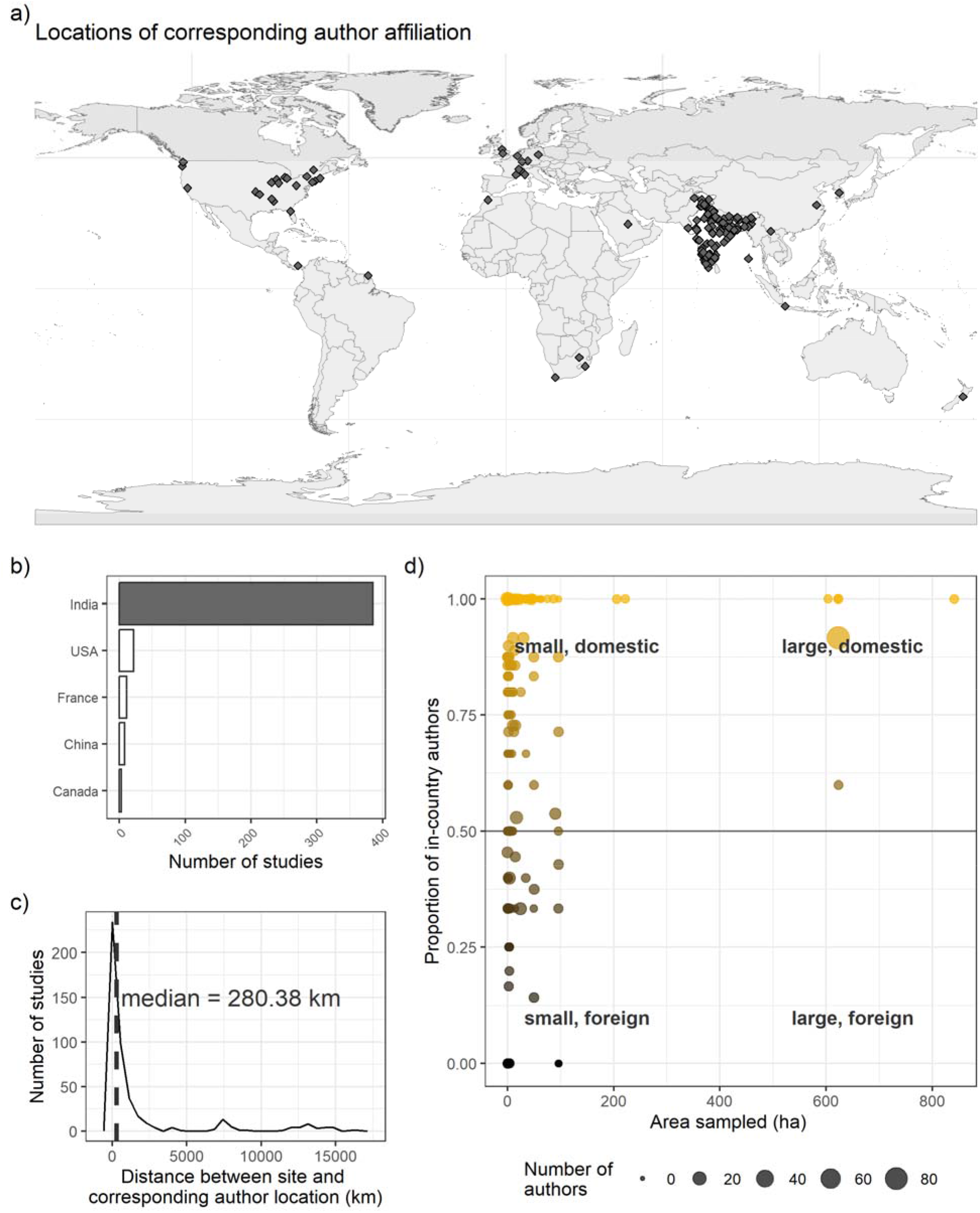
INvenTree authorship patterns and data equity. a) Locations of corresponding authors of papers in the INvenTree dataset, b) top five countries, representing 92.26%, where corresponding authors were affiliated; c) Measuring equity within the country - distance between corresponding author affiliation and study site across studies. d) Investigating parachute research - proportion of domestic and foreign authors in the INvenTree dataset and their relationship to area sampled. Size of the points represents number of authors in the study, colour is indicative of the proportion of in-country authors.

### Sampling priorities

The INvenTree dataset was used to identify sampling priority regions for ecological and conservation research (Fig 4). The INvenTree dataset showed disproportionate sampling across districts, with most (223 out of 341) districts with tree cover never sampled (Fig S6). When sampling gaps were visualised along tree cover and tree cover loss gradients, high heterogeneity among sites and biomes emerged (Fig 4). Notably, among districts with high tree cover, the northern Western Ghats, several districts of North-east India, parts of the Western Himalayas, and a few districts in eastern India fell in the lowest quantile of area sampled (Fig 4, Fig S6). Ecological sampling priority regions that fall in lower tree cover quantiles were many of the drier districts in peninsular India like the Deccan plateau, Eastern Ghats, Central India, as well as the western Himalayan foothills and the Nicobar Islands. There were also a few datasets reporting on tree-based inventories that fell outside of these forested districts (Fig S5). When considering sampling effort across ecoregions, the effort was highly skewed and most regions were highly undersampled relative to their extent (Fig S12).

**Figure 4:**
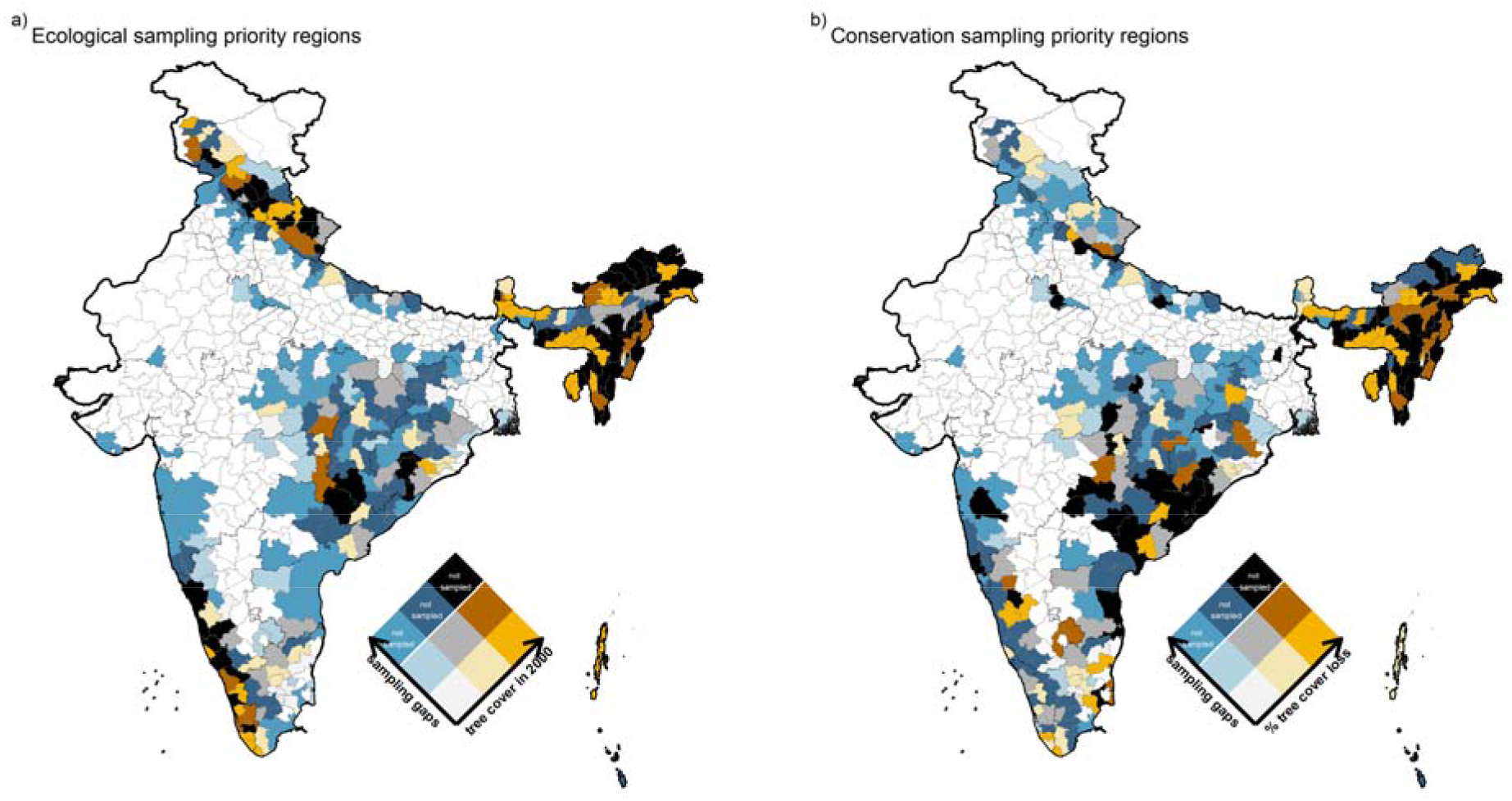
Ecological and conservation sampling priority regions. Bivariate chloropleth maps showing sampling gaps in plot-based tree inventories across districts in India in relation to a) mean tree cover in the district in 2000 and b) mean tree cover loss from 2000-2021 averaged over the area of the district. Sampling gaps were categorised into three categories; districts with 0 sampling effort were coded as the highest gap, the rest were split into two groups. Refer methods and supplementary for details.

Districts of the highest conservation sampling priority were largely different from those of the ecological sampling priority districts. Districts in the highest quantile of forest loss relative to forest cover were clustered mainly in north-east India and the eastern states of Odisha, Andhra Pradesh, Telengana and Chattisgarh (Fig S9). Among districts that fell in the highest quantile of tree cover loss, districts in the north-east and eastern India were the least studied (Fig 4). Data availability from areas of tree cover loss was poor from the northern Western Ghats, northeast India and the Nicobar Islands.

## Discussion

The INvenTree dataset, to our knowledge, is the first and largest meta-dataset of tree inventory information from India. This dataset covers all tree-based biomes of India and spans a total of 4653.64 ha. Using this dataset, we identify geographic and biome gaps in sampling and define ecological and conservation sampling priority regions. Most studies in the INvenTree dataset are authored by researchers affiliated to Indian universities and study local ecosystems. Although 80% of the dataset is not accessible, the strong presence of researchers in Indian organisations who are owners of high quality local data indicates potential for a regional network to promote a better understanding of ecological processes across larger spatial scales.

Harmonising these datasets under fair sharing protocols is a first step towards synthetic research in the region. Given India’s high biodiversity and human-dominated landscapes, such a network could promote a better understanding of ecosystems in the Anthropocene, while benefiting research and researchers.

Using the INvenTree dataset, we identified the extent and sampling biases across biomes and regions, providing directions to align future sampling of tree-based biomes within the country (Fig 4). The INvenTree dataset spans all of India’s diverse tree-based biomes, but shows non-uniform sampling across these. Among biomes, tropical and subtropical moist and dry forests were well sampled relative to their area. However, within India, there has been increased attention on the diversity of tropical dry forest types (Ratnam et al., 2016), their high biodiversity and threats to their conservation (Nerlekar et al., 2022, 2024). Our result highlights potential for these tree inventory data, once collated and standardized, to contribute to our larger understanding of dry tropical forests which remain among the least understood and most threatened tropical tree biomes globally (Miles et al., 2006; Siyum, 2020). Although the moist forests were better sampled, the distribution of the plots was clustered in the southern western ghats and north eastern forests, which are more speciose (Fig 1). Consequently, we report ecological sampling priority in the northern Western Ghats and north-east India and conservation sampling priority in north-east India as well as eastern India (Fig 4). Regional disparities in sampling can lead to biased understanding of ecological heterogeneity in patterns and processes which may result in inappropriate management, conservation or restoration practices. It is concerning that our proposed regions of priority conservation sampling (high loss/low sampling) overlap regions of large conservation concern with megaprojects (e.g. dams and oil palm expansion in North-East India, Bhattacharya (2022), Srinivasan et al. (2021); proposed development in the Nicobar Islands, Ramachandran (2022); mineral resource extraction in eastern India, Aggarwal (2021)). As these regions continue to lose tree cover, it is an urgent need to collect baseline information to support stakeholders in appropriate conservation action. Our proposed ecological and conservation sampling priority maps can provide a blueprint for the next decades of tree sampling in the country to help refine our ecological understanding and inform management and conservation.

### Challenges and promises of synthesising INvenTree data

A harmonised database of tree inventories from India, beyond the INvenTree meta-dataset, can fill crucial knowledge gaps in the human dimension of global tropical ecology to aid research, conservation and socio-cultural dimensions (Table 1). Large-scale datasets are more than the sum of their parts; for example, data published by Ramesh et al. (2010), covering the Western Ghats, has allowed for several syntheses, both at regional and global scales ranging from patterns and drivers of diversity to drivers of above ground carbon storage(e.g. Hardy et al. (2012); Krishnadas et al. (2021); Osuri et al. (2020); Davidar et al. (2018)). Spatially distributed datasets like the Forest Inventory and Analysis programme in the United States have led to at least 180 publications and increased the accuracy of predictive modeling of ecological communities (Tinkham et al., 2018). Similar to other national programmes, The Forest Survey of India (FSI) conducts country-wide tree inventories spanning ∼ 3500 ha total area every 5 years with a systematic gridded approach (https://fsi.nic.in/about-forest-inventory, Tewari (2016)). In comparison, studies in the INvenTree dataset in aggregate covers more than this area even after accounting for sites sampled multiple times. A harmonised database such as INvenTree will enable synthetic research on the patterns and processes structuring ecosystems across the region. Moreover, datasets with repeat sampling could also lend to ecological forecasting of ecosystems states with climate (e.g. Heilman et al. (2022)). Such data could immensely improve conservation efforts both by government and private actors by improving data available for monitoring and evaluation of forest management, restoration efforts, carbon credit projects and other nature-based solutions to mitigate climate change. These data would also be useful for universities and educators, making it possible to create data-driven educational products for ecology courses in India (e.g. Organisation for Tropical Studies courses in other geographies). We also envision that INvenTree data would serve as crucial baselines for practitioners and local communities involved in sustainable use of forest resources as well as “biocultural” conservation that balances conservation goals with local cultural values and worldviews (Sterling et al., 2017).

**Table 1.**
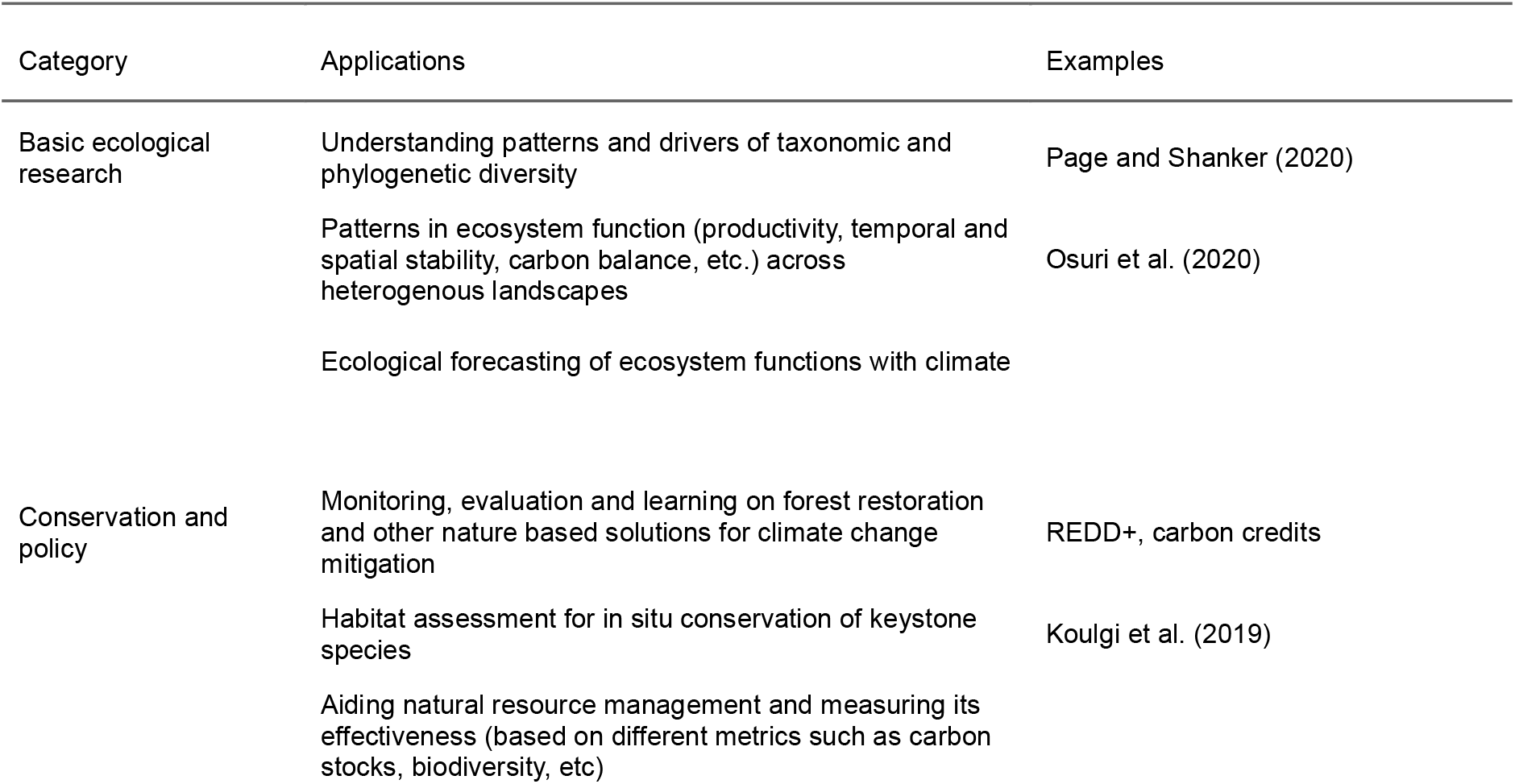
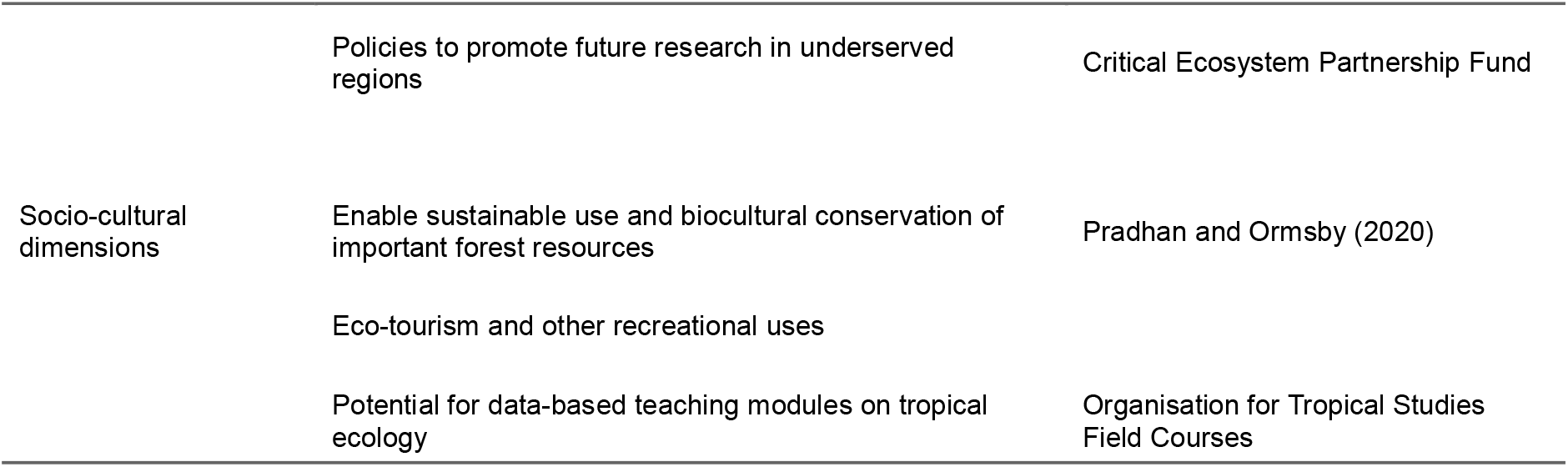
Potential applications emerging from a combined, harmonised INvenTree.

| Category | Applications | Examples |
| --- | --- | --- |
| Basic ecological research | Understanding patterns and drivers of taxonomic and phylogenetic diversity | Page and Shanker (2020) |
|  | Patterns in ecosystem function (productivity, temporal and spatial stability, carbon balance, etc.) across heterogeneous landscapes | Osuri et al. (2020) |
|  | Ecological forecasting of ecosystem functions with climate |  |
| Conservation and policy | Monitoring, evaluation and learning on forest restoration and other nature based solutions for climate change mitigation | REDD+, carbon credits |
|  | Habitat assessment for in situ conservation of keystone species | Koulgi et al. (2019) |
|  | Aiding natural resource management and measuring its effectiveness (based on different metrics such as carbon stocks, biodiversity, etc) |  |

| Category | Applications | Examples |
| --- | --- | --- |
|  | Policies to promote future research in underserved regions | Critical Ecosystem Partnership Fund |
| Socio-cultural dimensions | Enable sustainable use and biocultural conservation of important forest resources | Pradhan and Ormsby (2020) |
|  | Eco-tourism and other recreational uses |  |
|  | Potential for data-based teaching modules on tropical ecology | Organisation for Tropical Studies Field Courses |

The INvenTree dataset highlights the potential for collaborative approaches within the country to facilitate inclusion of high quality data into regional and global syntheses. Most research represented in the INvenTree dataset was conducted locally, likely integrating deep knowledge of landscape context, taxonomy and natural history. In heterogeneous tropical landscapes like India with high alpha and beta diversity, such localized, specialized data collection can be crucial for data quality. However, at present, the reusability of these data for further analyses is hindered by the small scale of individual datasets and their distributed ownership. Since many of these datasets have total sampling effort < 2 ha, the effort required to individually request permissions from authors and harmonise these may outweigh the benefits of their inclusion into previous global syntheses. An accepted set of best practices for data reusability is described by the FAIR principles for data; Findable, Accessible, Interoperable and Reusable (Wilkinson et al., 2016). Our assessment of the reusability of these datasets revealed that even while datasets were “findable” and “accessible”, the current forms in which they are publicly available most often did not meet reusability standards, especially for the finest grain, stem-level data, required for many syntheses. Despite the limited access at the stem level, plot-based vegetation studies are multi-layered in their data as well as further utility, allowing useful, scalable, synthetic research even with plot and species level data. Harmonising these heterogeneous datasets, to achieve “Interoperability” and “Reusability”, is difficult as they can be conducted using slightly different methodologies or adapted to context. Moreover, standardising species names across studies, including dealing with local names, requires taxonomic expertise and biogeographic knowledge (Singh, 2008). Assembling such a database would require close, continuous, collaborations with data owners and collectors.

The distributed inventories collated in INvenTree are important not just because of their extent, but the perspective of data sovereignty, and equity. FAIR principles for data sharing are not always fair for tropical forest data owners and making data unconditionally open could lead to further extractive science, as quantitative skills and publishing power tends to be concentrated in the global north and in elite universities (De Lima et al., 2022; Mori et al., 2015). Existing networks such as ForestPlots, respond to this inequity by insisting on co-authorship for primary data collectors (ForestPlots.net et al., 2021). We are encouraged by our findings that showcase a steady pattern of in-country authorship on tree-inventory-based publications from India, a positive contrast to the global pattern of parachute research in tropical landscapes (Asase et al., 2022; Miller et al., 2023). Acknowledging the enormous efforts and potential needs of current and future data owners, we envision a regional network that prioritises fair sharing policies as well as quantitative processing tools (e.g. open source code, tutorials, periodic data workshops). Combining frameworks for data sharing with tools and networks can empower Indian researchers, especially early-career researchers and those in under-funded universities to access and utilise these data for their research.

In conclusion, using the first synthesized metadataset of tree inventory data from India, we identified geographic gaps in sampling and highlighted priority regions for future sampling to fill critical gaps in tropical ecology. We also identify strong scholarship in forest ecology within India, demonstrated by a large number of authors and data owners affiliated to Indian institutions. Based on our findings, we propose The India Tree Inventory Network, a regional network of forest researchers from India that will promote synthetic forest research in India while prioritising data sovereignty and knowledge equity at the global and national scale. The INvenTree network will build on existing frameworks for data and knowledge sharing that acknowledge diverse contributions, costs and benefits inherent to an unequal world (Cooke et al., 2021; De Lima et al., 2022; Ocampo-Ariza et al., 2023).

### Positionality Statement

All authors of this paper are Indian nationals, working in ecology and allied fields, with substantial field experience in India. KA and NM are currently postdoctoral researchers in US institutions and were also affiliated with US universities for their PhD, but have studied and worked in Indian institutions up to the Masters level. KA has led field data collection in India through all stages of her professional life starting from undergrad. AG and AK are PhD students in Indian institutions. AS is affiliated with US institutions for his PhD and has studied and worked in India until the Masters level. AJ is a Masters student in Europe. TN is a PhD student in the UK doing field work in India and has studied in India until her undergraduate degree. SO is a PhD student in Germany. MS is a Professor at an Indian institution.

## Acknowledgements

We are grateful to the community of data collectors and owners contributing to forest ecology in India. KA acknowledges the support of the Smithsonian Institution Postdoctoral Fellowship during the completion of this manuscript.

## Data Accessibility Statement

The data and code used in this manuscript will be made available on the India-INvenTree GitHub repository (www.github.com/India-INvenTree/metadata-analysis) and on Zenodo upon acceptance of this manuscript.

