## Supplementary figures and tables for "Harmonising distributed tree inventory datasets across India can fill critical gaps in tropical ecology"

### Supplementary Information: Harmonising distributed tree inventories across India can fill critical gaps in tropical ecology (Anujan et al., *TBD*)

#### Appendix S1. INvenTree data spread on other base layers

We used the Rodgers & Panwar (1988) classification of biogeographic zones to quantify data spread across the 10 different zones in India. This map classifies India into zones based on biogeographic history of the region, each of which may have multiple biomes structured by distinct ecological processes. On the other hand, biome classifications could have uncertainties, especially in their boundaries and transition zones. Since this classification does not assume biome boundaries based on climate etc, it complements the Dinerstein et al. (2017) biome and ecoregion classification for the region.

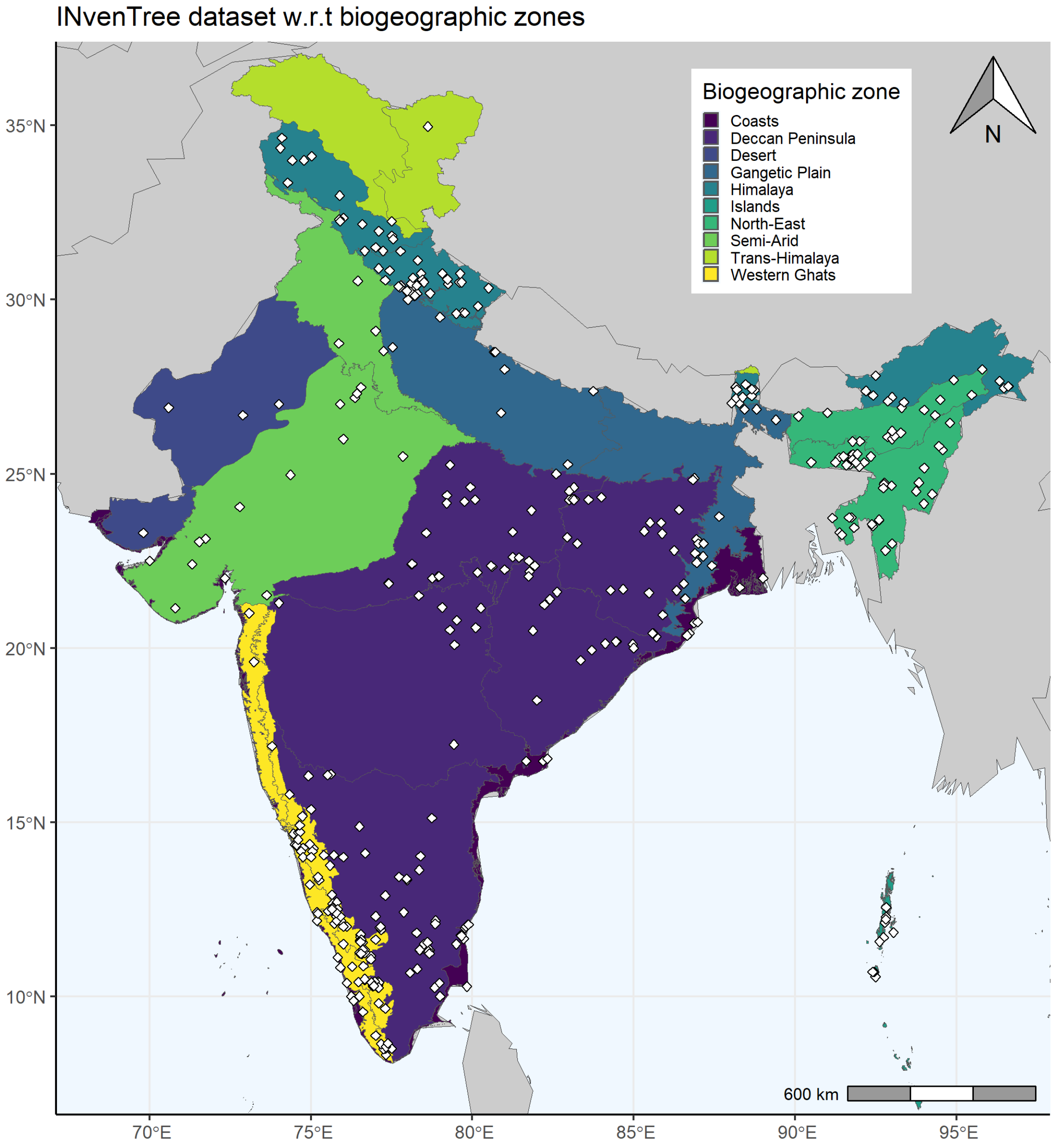

Figure S1. INvenTree data spread with respect to biogeographic zones in India.

We also looked at data spread across ecoregions defined by Dinerstein et al. (2017)

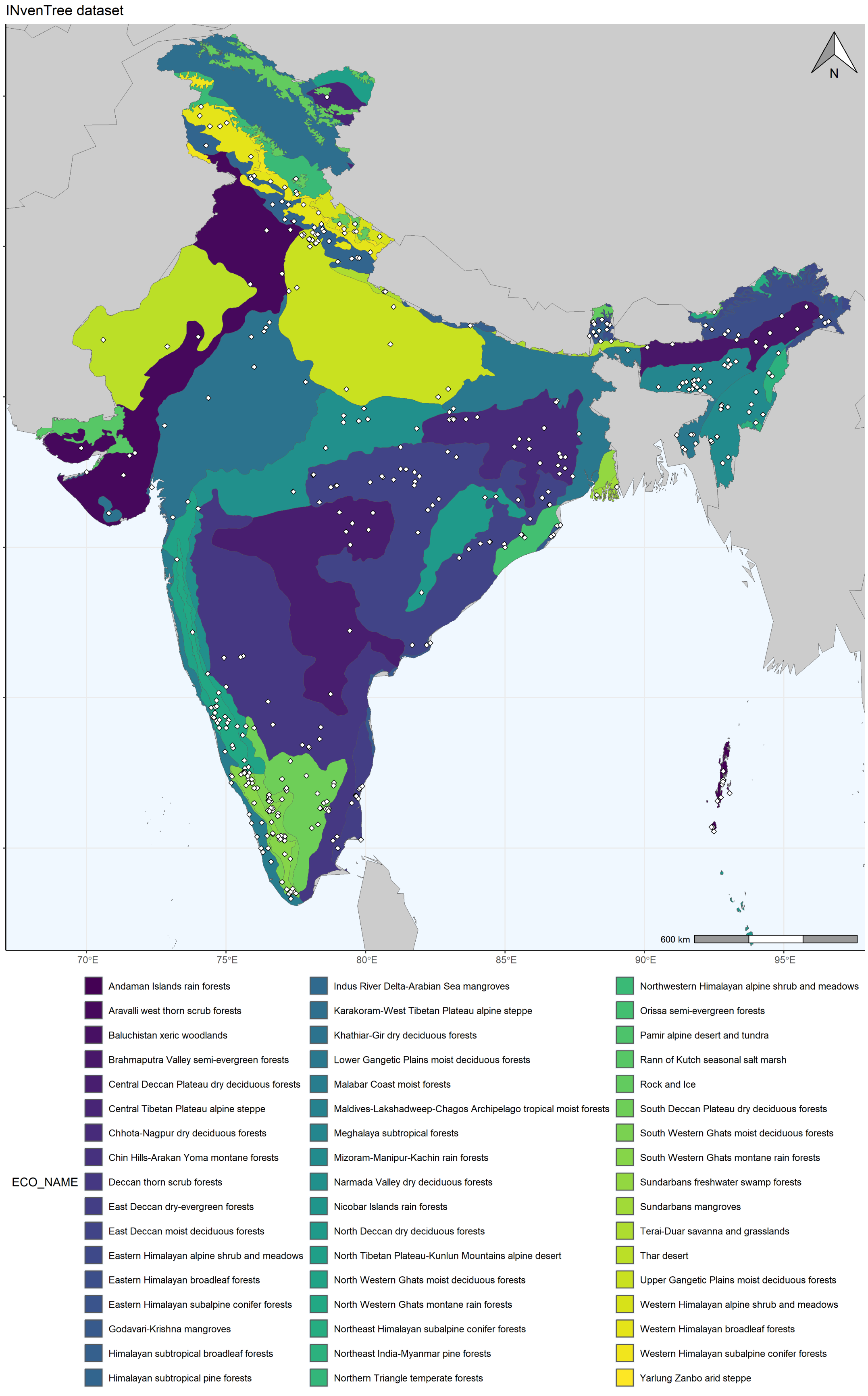
INvenTree dataset
35°N
30°N
25°N
20°N
15°N
10°N
70°E
75°E
80°E
85°E
90°E
95°E
N
600 km
ECO_NAME
Andaman Islands rain forests
Aravalli west thorn scrub forests
Baluchistan xeric woodlands
Brahmaputra Valley semi-evergreen forests
Central Deccan Plateau dry deciduous forests
Central Tibetan Plateau alpine steppe
Chhota-Nagpur dry deciduous forests
Chin Hills-Arakan Yoma montane forests
Deccan thorn scrub forests
East Deccan dry-evergreen forests
East Deccan moist deciduous forests
Eastern Himalayan alpine shrub and meadows
Eastern Himalayan broadleaf forests
Eastern Himalayan subalpine conifer forests
Godavari-Krishna mangroves
Himalayan subtropical broadleaf forests
Himalayan subtropical pine forests
Indus River Delta-Arabian Sea mangroves
Karakoram-West Tibetan Plateau alpine steppe
Khathiar-Gir dry deciduous forests
Lower Gangetic Plains moist deciduous forests
Malabar Coast moist forests
Maldives-Lakshadweep-Chagos Archipelago tropical moist forests
Meghalaya subtropical forests
Mizoram-Manipur-Kachin rain forests
Narmada Valley dry deciduous forests
Nicobar Islands rain forests
North Deccan dry deciduous forests
North Tibetan Plateau-Kunlun Mountains alpine desert
North Western Ghats moist deciduous forests
North Western Ghats montane rain forests
Northeast Himalayan subalpine conifer forests
Northeast India-Myanmar pine forests
Northern Triangle temperate forests
Northwestern Himalayan alpine shrub and meadows
Orissa semi-evergreen forests
Pamir alpine desert and tundra
Rann of Kutch seasonal salt marsh
Rock and Ice
South Deccan Plateau dry deciduous forests
South Western Ghats moist deciduous forests
South Western Ghats montane rain forests
Sundarbans freshwater swamp forests
Sundarbans mangroves
Terai-Duar savanna and grasslands
Thar desert
Upper Gangetic Plains moist deciduous forests
Western Himalayan alpine shrub and meadows
Western Himalayan broadleaf forests
Western Himalayan subalpine conifer forests
Yarlung Zanbo arid steppe

Figure S2. INvenTree data spread with respect to ecoregions in India. Ecoregions are classified based on the ecoregion classification by Dinerstein et al. (2017)

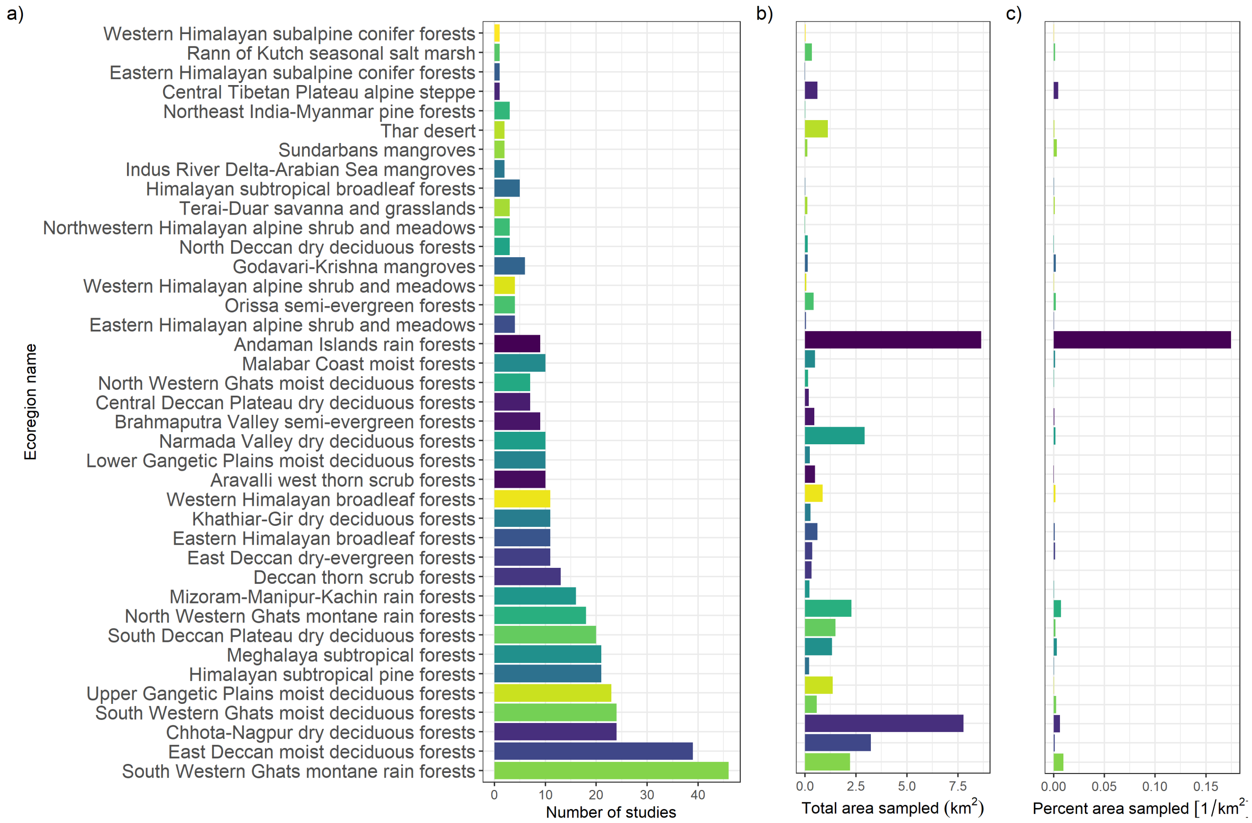

Figure S3. INvenTree dataset distribution across ecoregions in number of studies, area and proportion sampled

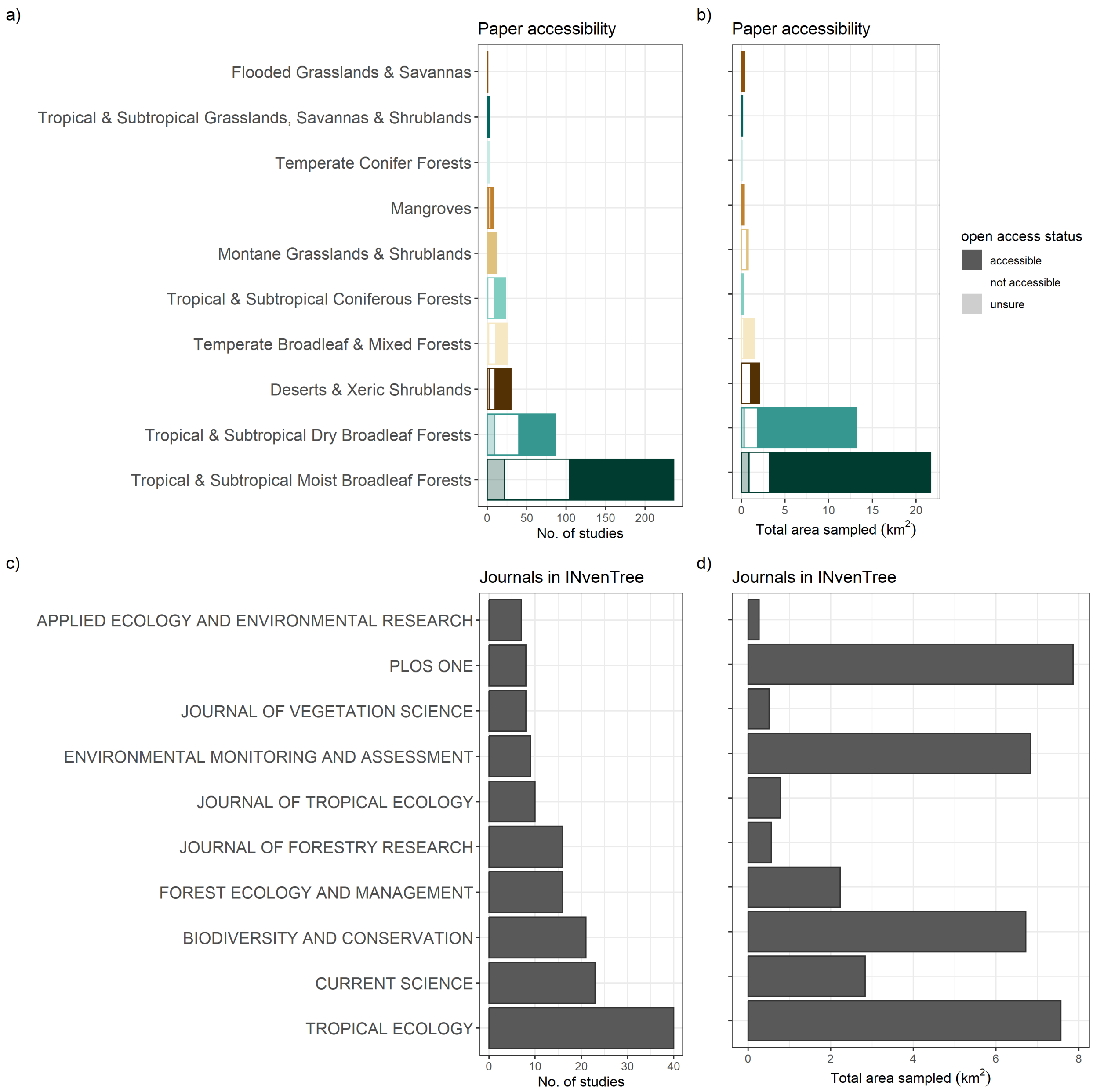

Figure S4. Open access status across the INvenTree metadataset and journal composition

#### Appendix S2. Records of studies where corrections were applied

For all studies, we uses the geographic coordinates of the study locations and extracted the district ID and biome from corresponding shapefiles. However, for some studies in the INvenTree dataset, the coordinates did not result in appropriate district ID or biomes. Most often this was because of small mismatches in the boundaries of shapefiles, especially for border territories or coastal ecosystems.
Below is a record of such studies where we manually assigned a biome or district based on the information in the paper.

Table S1. Studies with missing districts and manually assigned districts.

| districtID | paperID | DOI | State | District |
|---|---|---|---|---|
| 1 | A1855 | https://doi.org/10.1007/s11852-015-0398-4 | Andaman and Nicobar | Andaman Islands |
| 118 | A0794 | https://doi.org/10.1016/j.heliyon.2020.e04685 | Gujarat | Bharuch |
| 178 | C0221 | link | Jammu and Kashmir | Ladakh (Leh) |
| 378 | A1701 | https://doi.org/10.18520/cs/v110/i12/2253-2260 | Orissa | Kendrapara |
| 491 | C0029 | 10.4236/oje.2016.610057 | Tripura | South Tripura |
| 492 | A0505 | https://doi.org/10.1007/s11270-021-05133-z | Tripura | West Tripura |
| 580 | A1335 | https://doi.org/10.1016/j.japb.2018.01.012 | West Bengal | Darjiling |
| 580 | A0228 | https://doi.org/10.1038/s41598-022-08483-8 | West Bengal | Darjiling |

Table S2. Studies with missing biomes and manually assigned values.

| paperID | DOI | Biome | Ecoregion |
|---|---|---|---|
| A0447 | https://doi.org/10.1007/s10750-021-04651-5 | Mangroves | Andaman Islands rain forests |
| A1212 | https://doi.org/10.14719/pst.2019.6.2.435 | Tropical & Subtropical Moist Broadleaf Forests | Brahmaputra Valley semi-evergreen forests |
| A1524 | https://doi.org/10.1007/s10531-017-1344-6 | Tropical & Subtropical Dry Broadleaf Forests | Central Deccan Plateau dry deciduous forests |
| A0809 | https://doi.org/10.1371/journal.pone.0235783 | Deserts & Xeric Shrublands | Deccan thorn scrub forests |
| A1539 | NA | Temperate Conifer Forests | Northeast India-Myanmar pine forests |
| A1850 | link | Temperate Broadleaf & Mixed Forests | Himalayan subtropical broadleaf forests |
| A1158 | https://doi.org/10.1007/s11676-018-0600-2 | Mangroves | Malabar Coast moist forests |
| A0365 | https://doi.org/10.3389/fenvs.2021.724950 | Tropical & Subtropical Moist Broadleaf Forests | Meghalaya subtropical forests |
| A0279 | https://doi.org/10.1111/btp.13068 | Tropical & Subtropical Dry Broadleaf Forests | Narmada Valley dry deciduous forests |
| A1722 | no DOI (manual) | Tropical & Subtropical Dry Broadleaf Forests | Narmada Valley dry deciduous forests |
| C0388 | https://doi.org/10.1890/10-0133.1 | Tropical & Subtropical Moist Broadleaf Forests | North Western Ghats montane rain forests |
| A2104 | https://doi.org/10.5194/isprsarchives-XL-8-651-2014 | Tropical & Subtropical Moist Broadleaf Forests | Orissa semi-evergreen forests |
| A0704 | https://doi.org/10.18520/cs/v119/i9/1517-1525 | Flooded Grasslands & Savannas | Rann of Kutch seasonal salt marsh |
| A2585 | https://doi.org/10.1016/j.flora.2010.04.011 | Tropical & Subtropical Dry Broadleaf Forests | South Deccan Plateau dry deciduous forests |
| A1446 | https://doi.org/10.1111/jvs.12586 | Tropical & Subtropical Dry Broadleaf Forests | South Deccan Plateau dry deciduous forests |
| A2056 | https://doi.org/10.1007/s11676-013-0396-z | Tropical & Subtropical Moist Broadleaf Forests | Upper Gangetic Plains moist deciduous forests |
| A1335 | https://doi.org/10.1016/j.japb.2018.01.012 | Temperate Broadleaf & Mixed Forests | Himalayan subtropical broadleaf forests |
| A0794 | https://doi.org/10.1016/j.heliyon.2020.e04685 | Mangroves | Indus River Delta-Arabian Sea mangroves |
| C0010 | no DOI (manual) | Mangroves | Sundarbans mangroves |
| A2965 | https://www.jstor.org/stable/pdf/24109319.pdf | Tropical & Subtropical Moist Broadleaf Forests | Andaman Islands rain forests |
| A0228 | https://doi.org/10.1038/s41598-022-08483-8 | Temperate Broadleaf & Mixed Forests | Himalayan subtropical broadleaf forests |
| A1115 | https://doi.org/10.2112/SI86-031.1 | Mangroves | Indus River Delta-Arabian Sea mangroves |
| A1855 | https://doi.org/10.1007/s11852-015-0398-4 | Mangroves | NA |
| A1701 | https://doi.org/10.18520/cs/v110/i12/2253-2260 | Mangroves | Godavari-Krishna mangroves |

#### Appendix S3. Additional methods for analysing FAIR principles in INvenTree metadataset

To analyse the reproducibility of the published datasets, we used the FAIR principles (Wilkinson et al., 2016) as a framework. This framework classifies the reproducibility of datasets in 4 steps - “Findable”, “Accessible”, “Interoperable” and “Reproducible”. During manual sorting of datasets, we noted whether data was “Findable” in some form beyond broad summaries. For the ~25% of datasets where the data was “findable”, we categorised the “FAIR”ness of the data for three different levels of graininess: plot-level data, species-level data and stem-level data. These independent axes, of graininess and access, allowed us to get a practical understanding of the true utility of these data as they stand, as well as the immediate needs to make them more useful for future research.

Our categories for graininess included:
-Plot-level data: Data that is aggregated at the plot level, with “plot” being the smallest unit of data collection.
-Species-level data: Data that is aggregated at the species level. This may or may not be aggregated at the plot level.
-Stem-level data: Data at the individual stem level, with separate attributes, like DBH, reported for each stem.

For each paper, under each of these three categories, we noted the following:
- *“Findable”*:
— data has separate DOI
— data is reported (e.g. plot-level summaries)
-*“Accessible”*:
— metadata is available (plot lat/long for plot-level, species names for species-level, locations for stems)
— data is available in a machine-readable format (csv or pdf)
-*“Interoperable”*:
— data is available in a standard format (e.g. species scientific names are reported)
— data is available in a standard unit (e.g. DBH in cm)
-*“Reproducible”*:
— data has a license

#### Appendix S4. Additional methods for sampling priority maps

We first used the Hansen et al. (2013) dataset to find areas of forest cover and forest cover loss in India. This dataset is at the 30 m x 30 m resolution corresponding to LANDSAT imagery. We used the MODIS LC criterion of tree cover >30% to be tree-based biomes (Friedl & Sulla-Menashe, 2022) on the tree cover layer to identify pixels that had tree cover in the year 2000, in the middle of our period of interest. This allowed us to include woody savannas in the analysis and we excluded savannas that were predominantly grass/herbaceous (usually categorised as <30% tree cover). We also used the Hansen et al. (2013) tree cover loss layer to identify pixels that experienced stand-clearing disturbances in the time period between 2000 and 2012. We visualise INvenTree data spread relative to these two layers here.

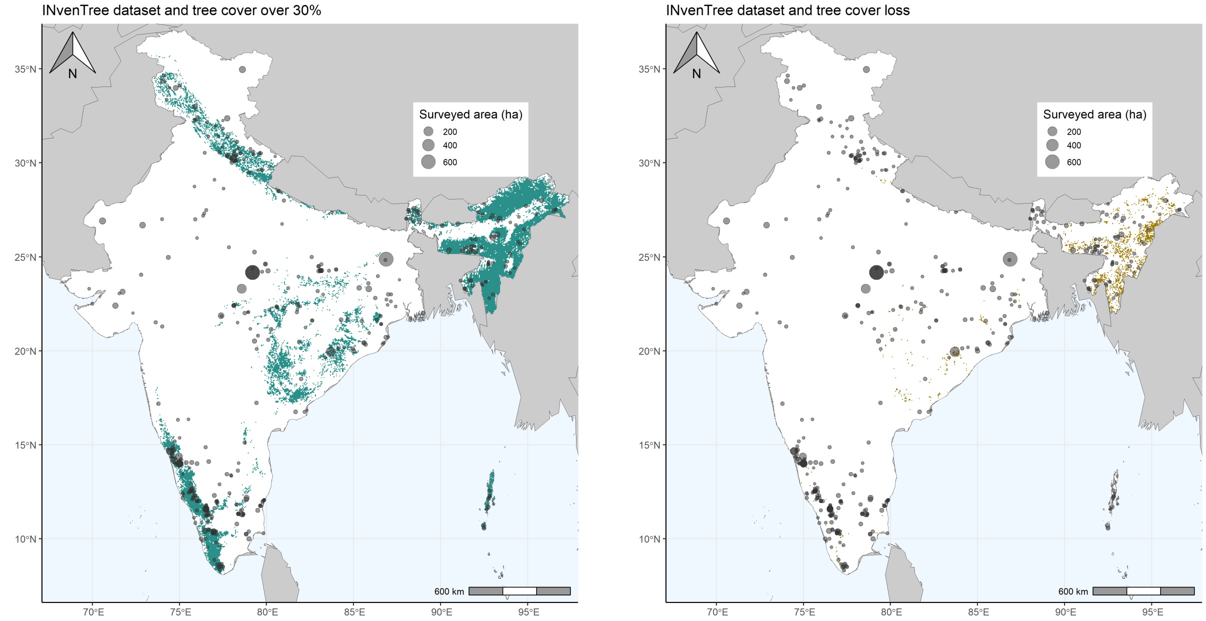

Figure S5. INvenTree sampling effort across forest cover and loss

We then identified 314 districts with non-zero forest cover in 2000 (using the above delimitation). Although the INvenTree dataset had sampling outside of these areas, we chose to restrict our sampling priority analyses to only regions of high tree cover, as there are likely to be more local patterns and high uncertainties outside of these. For each forested district, we calculated the percent of forest area, percent of total area experiencing forest loss, percent of forested area that was lost, percent of total area sampled and percent of forested area sampled. Along each of these variables, we divided the district into tertiles (0-33%, 33-66% and 66-100%). Forest loss and sampling variables had high skew with more than half the districts having 0 values. Moreover, in both of these cases, the 0 category needs to considered seriously and separate from the smallest tertile. Therefore, for sampled area and loss, we assigned the districts with 0 sampling (or loss) to one category and evenly split the remaining.

The tertiles of each of these variables are individually visualised below. Where there is large skew, histograms are shown on a log scale.

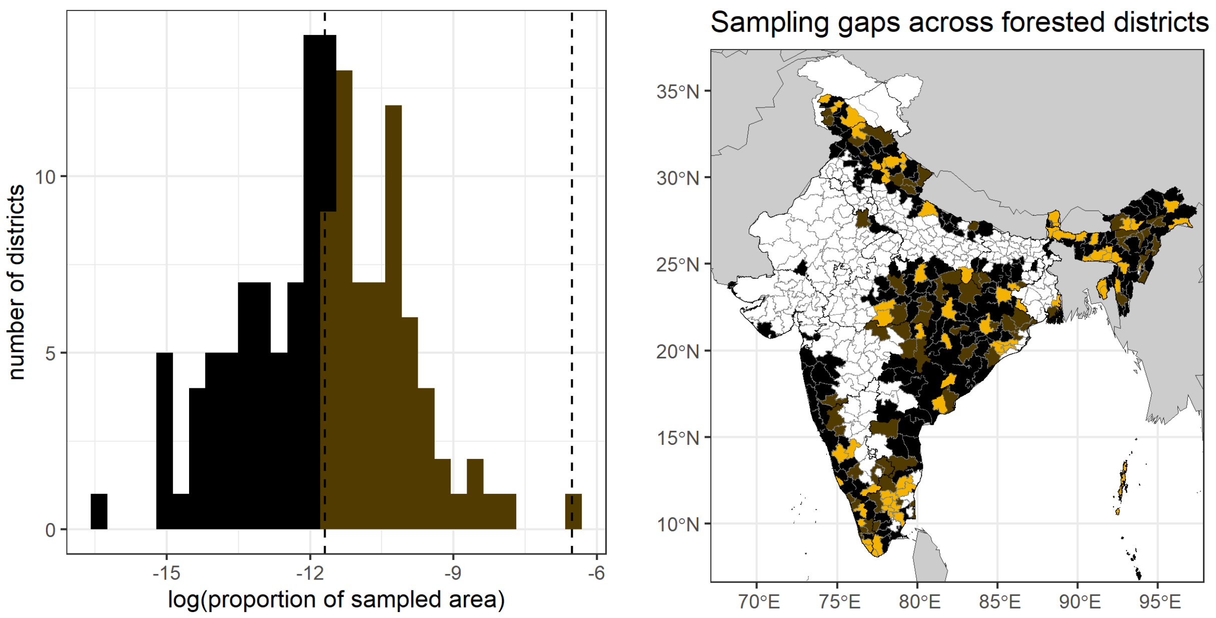

Figure S6. Sampling gaps across districts divided into tertiles. All districts with 0 sampling have been assigned highest value of gap, rest equally divided

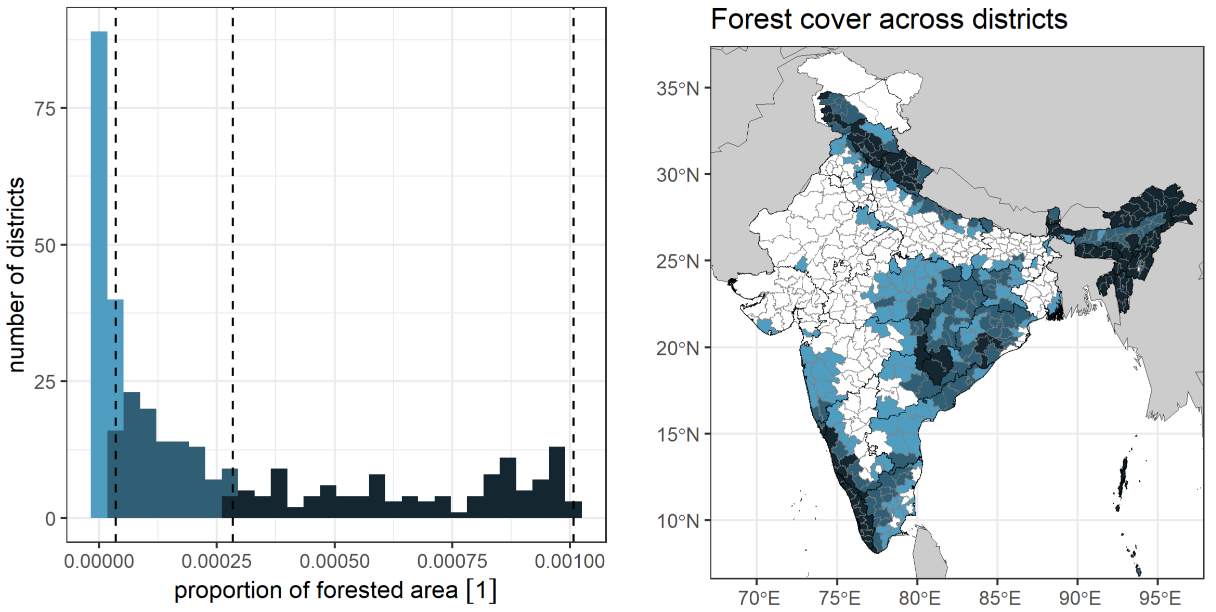

Figure S7. Forest cover across forested districts divided into tertiles.

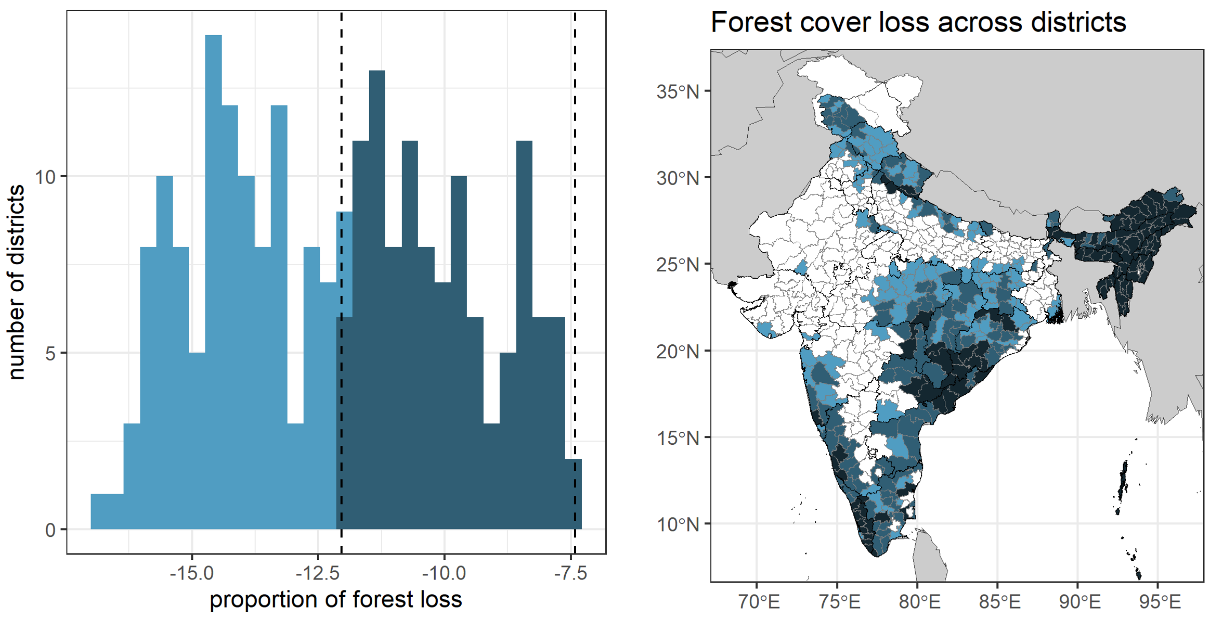

Figure S8. Absolute forest cover loss across forested districts divided into tertiles.

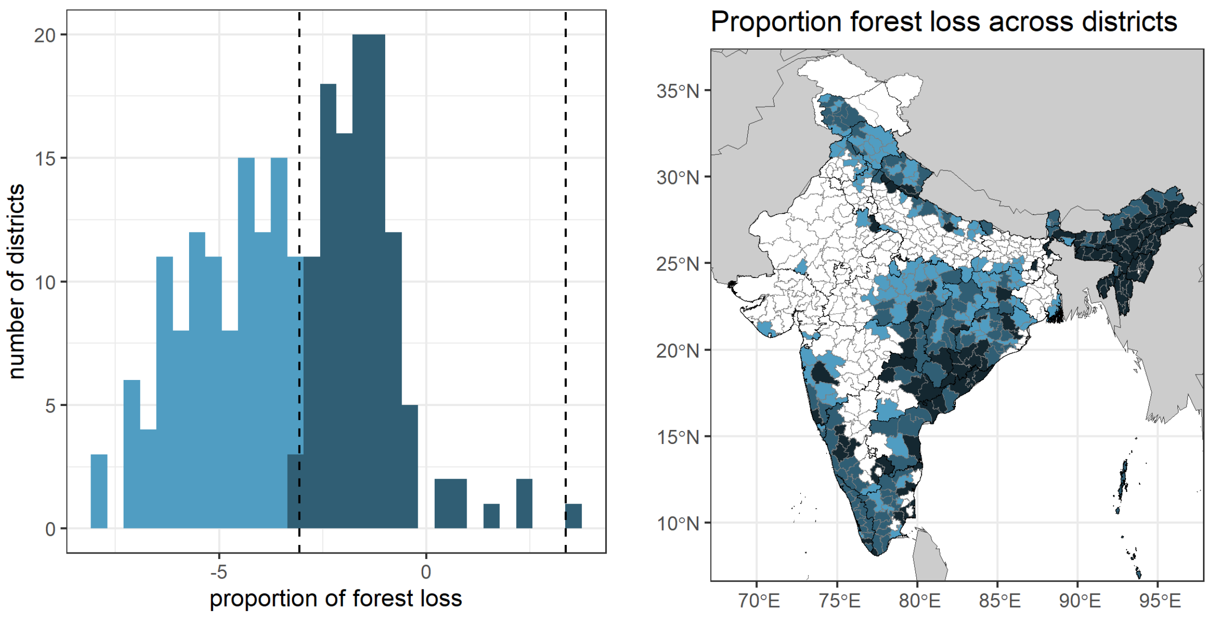

Figure S9. Forest cover loss relative to forest cover across forested districts divided into tertiles.

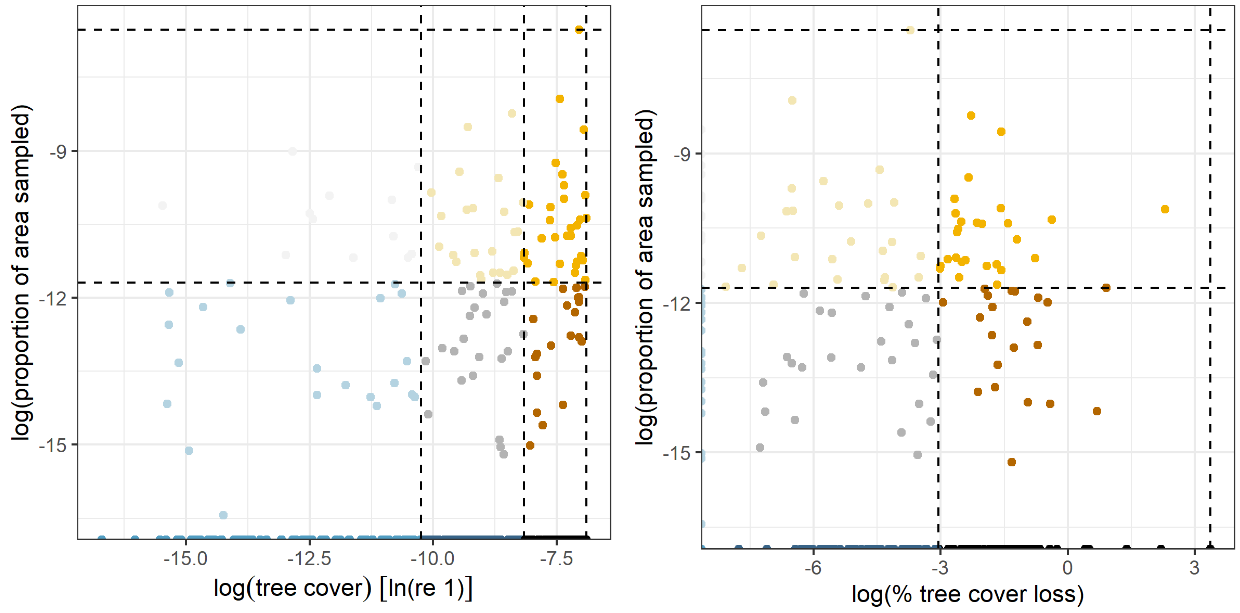

Figure S10. Distribution of districts along bivariate axes of sampling gap and forest cover or loss

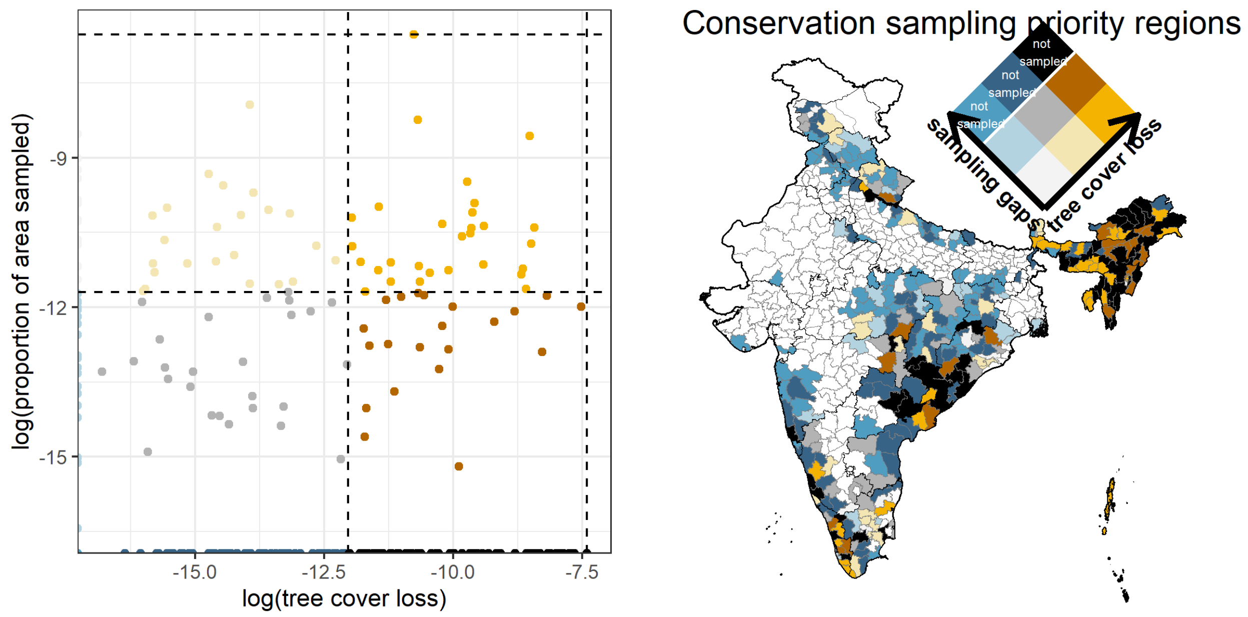

Figure S11. Data distribution of absolute forest loss and proportion of area sampled and the associated bivariate chloropleth map

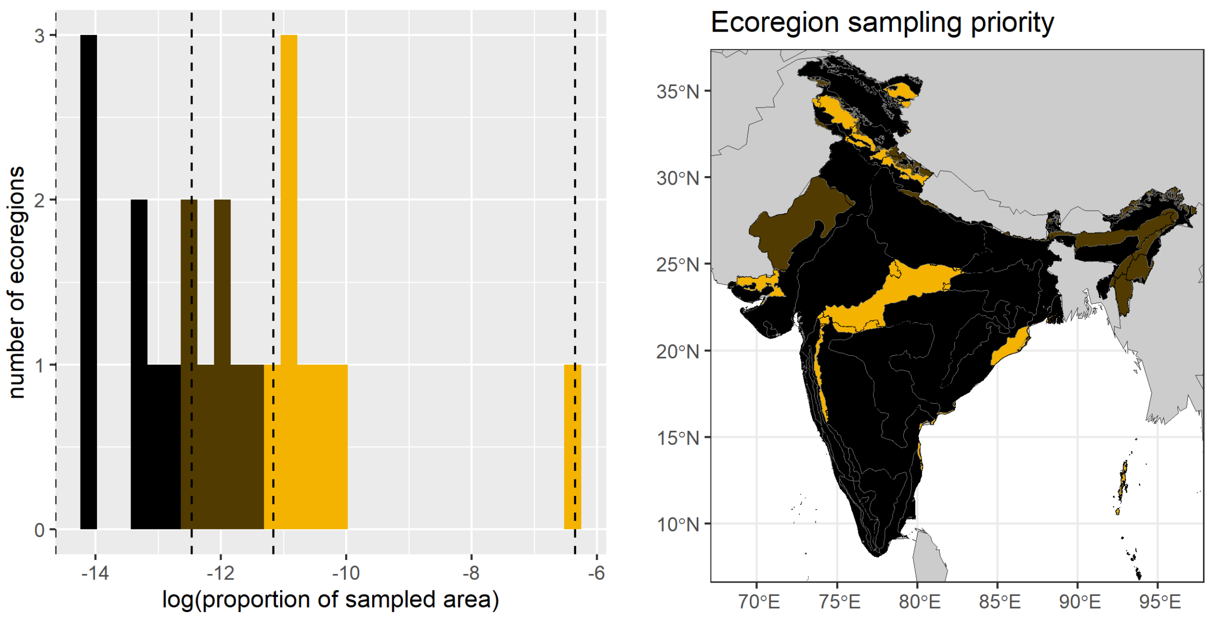

Figure S12. Sampling gaps across ecoregions

#### Appendix S5. Supplementary results for authorship analysis

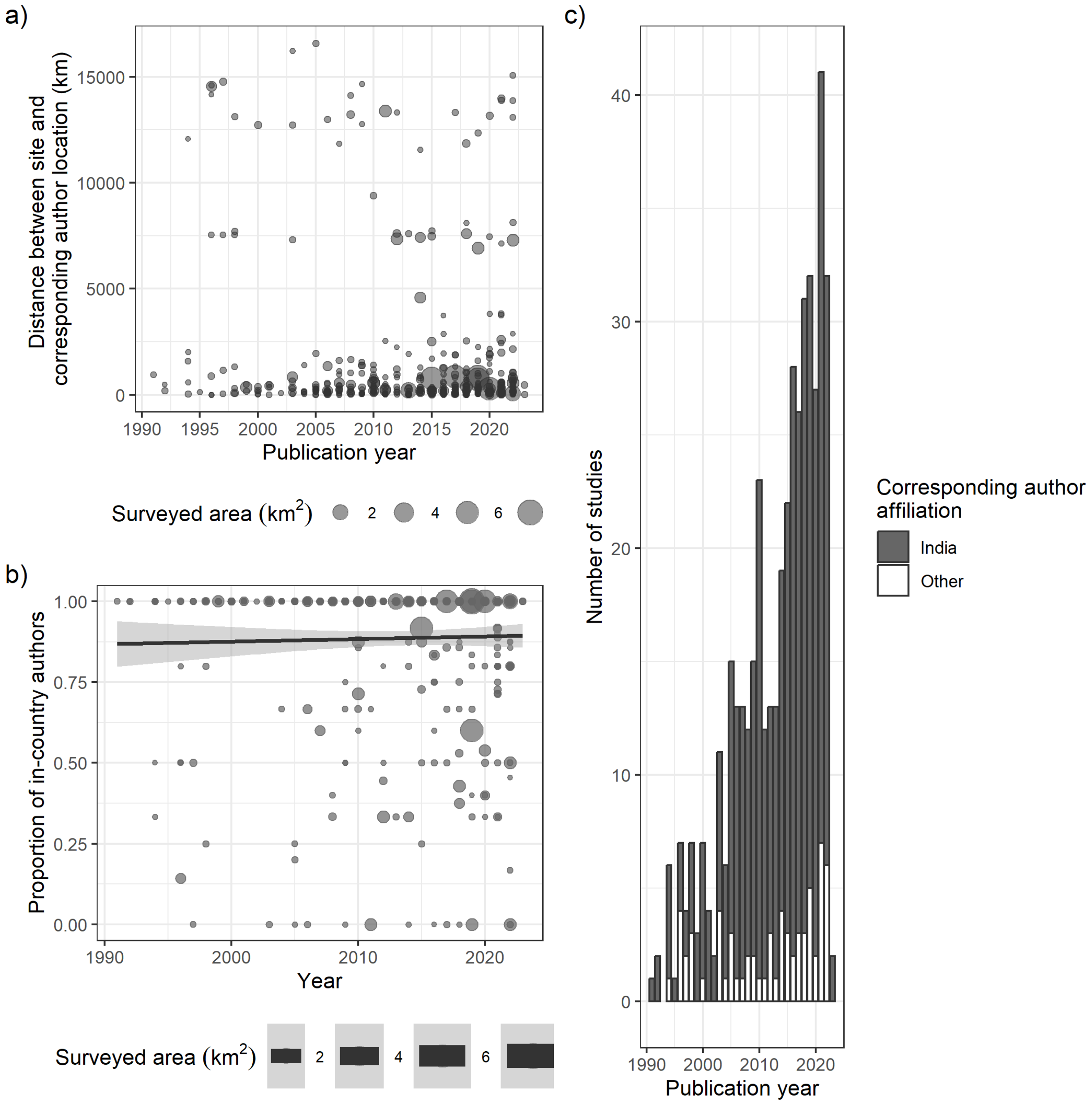

Figure S13. Authorship across time
